# Standing genetic variation and *de novo* mutations underlie parallel evolution of island bird phenotypes

**DOI:** 10.1101/2023.06.29.546893

**Authors:** Andrea Estandía, Ashley T. Sendell-Price, Bruce C. Robertson, Sonya M. Clegg

## Abstract

Parallel evolution occurs when the same trait evolves in closely related lineages in response to similar ecological contexts and provides some of the best examples of determinism in evolutionary biology. However, from a genetic standpoint, this process can be driven by either new mutations that appear independently in each diverging population or by selection on existing genetic variation common to both lineages. Small-bodied birds, for example, tend to increase in size after they colonise a new island, following what is known as the ‘island rule’. Such is the case of the silvereye (*Zosterops lateralis*), a prolific natural coloniser of southwest Pacific islands. Island forms of this bird species increase in body size after they establish, with the pattern and pace of change consistent with directional natural selection and evident even in the most recent colonisations within the last 200 years. The system provides an exceptional opportunity to explore the genomic basis of repeated body size evolution. We sequenced 377 whole genomes from 31 different silvereye populations, which revealed that both mechanisms are at play: in some lineages, new mutations are highly associated with body and bill size, but there are also highly associated polymorphisms present across all populations. Our research sheds light on the genomic basis of repeated body size evolution and emphasises that multiple molecular mechanisms can underlie similar evolutionary trajectories even within a single taxon.

## Introduction

Parallel evolution refers to the independent emergence of similar traits in different lineages that share a common ancestor (Cerca 2023), and illustrates deterministic evolutionary outcomes that are widespread across the Tree of Life (Blount et al. 2018). Natural selection is typically responsible for parallel evolution, as genetic drift is unlikely to produce such similar changes (Arendt and Reznick 2008; Stayton 2008; Losos 2011; McGhee 2011). Selection can act on standing genetic variation to produce the same phenotype (Lai et al. 2019; Therkildsen et al. 2019; Konečná et al. 2021), on different *de novo* mutations in each population (Fulgione et al. 2022; Cerca et al. 2023), or a combination of the two (Ament-Velásquez et al. 2022). However, a faster rate of adaptive change is more often facilitated from standing variation than from mutations as the latter are mostly neutral or mildly deleterious (Hedrick 1983; Barrett and Schluter 2008). For example, the rapid repeated evolution of stickleback low-plated phenotypes is driven by reassembling ancestral variation (Colosimo et al. 2005; Archambeault et al. 2020). Additionally, standing genetic variation can be supplemented with genetic variants that contribute to parallel evolution shared via admixture among recently diverged lineages (Taylor et al. 2020; Montejo-Kovacevich et al. 2022; Cerca et al. 2023; Wang et al. 2023). The same processes can affect single polymorphisms or full blocks in the genome: chromosomal rearrangements, like inversions, can play a role in parallel evolution by keeping sets of adaptive alleles together in strong linkage disequilibrium (Faria et al. 2019; Harringmeyer and Hoekstra 2022). Understanding the relative importance of alternative genetic mechanisms behind parallel evolution has received considerable interest over the last few decades (MacPherson and Nuismer 2017; Westram et al. 2022). Genomic methods now allow us to investigate examples spanning diverse systems, providing empirical data that is necessary to draw generalities.

Islands provide environments where comparable evolutionary pressures can result in the rapid emergence of similar phenotypes (Baeckens and Van Damme 2020). The ‘island syndrome’ refers to the phenomenon where sets of morphological, physiological, and behavioural traits evolve in a predictable way when organisms colonise islands. The syndrome ultimately results from the exposure to a common suite of biotic conditions on islands; a combination of reduced predation and parasitism, and a shift in the balance of intra- and interspecific competition, features that fundamentally change the selective pressures in predictable and repeatable ways (Clegg and Owens 2002; Meiri et al. 2004; Lomolino 2005). Arguably the best-characterised component of this syndrome is the ‘island rule’ that describes evolution towards medium body sizes in insular settings; small colonising species evolve larger body sizes and vice versa (Foster 1964; Clegg and Owens 2002; Benítez-López et al. 2021). There is evidence for the island rule across vertebrate taxa, including birds (Benítez-López et al. 2021). Bill size also follows this rule whereby short-billed species show an increase in size (Clegg and Owens 2002). Body and bill size are complex multivariate quantitative traits predicted to be highly polygenic (Yang et al. 2011b; Bosse et al. 2017; Duntsch et al. 2020), but some studies in natural populations show that a few genes of major effect can also impact these traits (Lamichhaney et al. 2015, 2016; vonHoldt et al. 2018; Duntsch et al. 2020).

Members of the silvereye subspecies complex (*Zosterops lateralis*, Aves) have repeatedly colonised islands across the South Pacific from the Australian mainland and Tasmania, with up to 17 morphological subspecies currently recognised (Clements et al. 2021). Insular silvereyes are larger than mainland ones, and the degree of this shift increases with time since colonisation (Clegg et al. 2002a), a pattern that has been shown to be a consequence of directional selection rather than drift (Clegg and Owens 2002; Clegg et al. 2008). The island populations represent at least four independent colonisations from the Australian mainland: to Southern Melanesia (∼ 150,000 years ago), Lord Howe Island (∼ 100,000 years ago), southern Great Barrier Reef Islands, including Heron Island (∼ 4000 years ago) and a recent historically recorded sequential colonisation sequence from Tasmania to New Zealand and outlying islands (<200 years ago) (Black 2010) (Fig. 1). The populations arising from these independent colonisations are genetically isolated from the original mainland source and other colonisations, though gene flow occurs among Southern Melanesian islands and among recently colonised populations (Estandía et al. 2023). The combination of independent and known colonisation sequences, along with broadly repeated shifts in body and bill size in similar ecological contexts provides an excellent system to explore the genetic mechanisms of parallel evolution. Here, we ask whether *de novo* mutations or standing genetic variation lead to the same phenotype: increased body and bill size in small-bodied species after island colonisation. We find that both are associated with body size variation in Southern Melanesia, whereas only standing genetic variation appears important for repeated patterns in all other populations. This demonstrates empirically that two genetic mechanisms hypothesised to underlie repeated evolution can operate simultaneously in a single system.

**Fig. 1.**
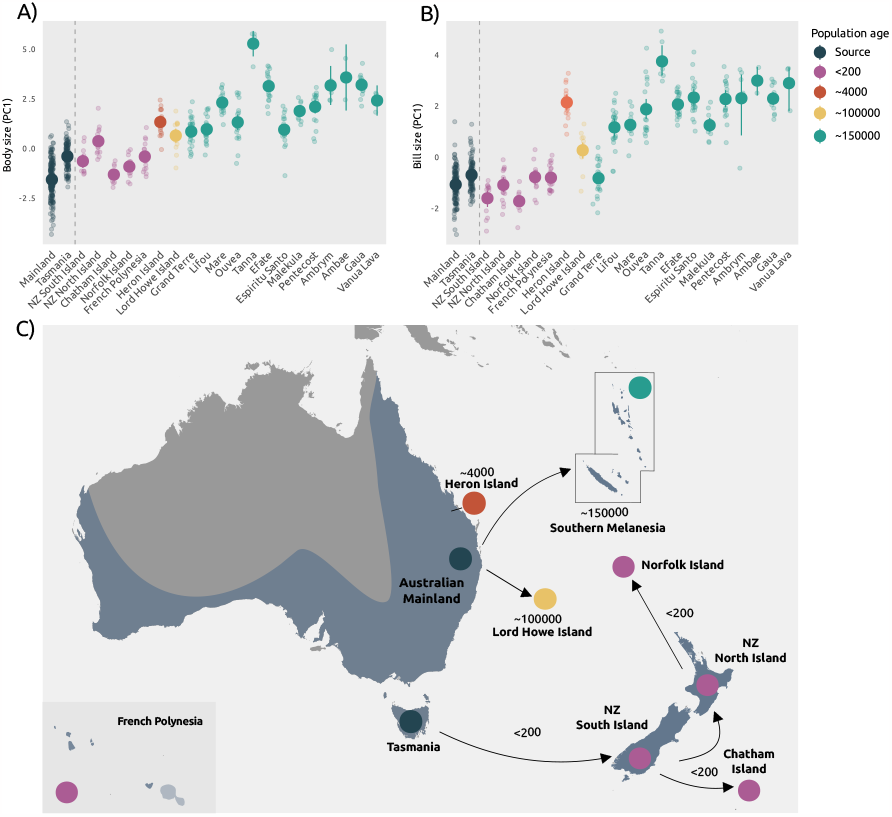
Morphological divergence in A) body and B) bill size (PC1) among silvereye forms. The source populations are placed left to the vertical dashed line. Colonisations are arranged chronologically: the most recent ones on the left and the most ancient on the right. C) The map shows the silvereye distribution highlighted in blue and the time of colonisation of each island or archipelago. French Polynesia is shown as inset.

## Results

### The genetic basis of repeated body and bill size evolution

To describe and quantify the bill and body size shifts in the populations studied here, we summarised univariate traits as principal component scores, resulting in a PC1 for body, and a PC1 for bill that represented size. In both cases 75% of the variation was explained by PC1 and roughly equal trait loadings of univariate traits (Fig. 1; Table S1). We statistically confirmed increases in body and bill size (Fig. S1; Table S2). To examine the genetic basis of repeated evolution, we generated low-to-intermediate coverage (5X) whole-genome sequencing data of 377 silvereyes from 24 islands and sampled across seven Australian States and Territories representative of most of the silvereye range (Table S3). After alignment of the sequenced individuals, we analysed genetic variation in a probabilistic framework accounting for low coverage (ANGSD (Korneliussen et al. 2014)) and obtained 14.3 million SNPs with minor allelic frequencies higher than 5% for differentiation analyses.

Population structure was described by a genomic PCA based on whole-genome data that revealed three major groups (Fig. S2 A). PC1 explained 20.18% of the total variance and separated two clusters: one containing individuals from Southern Melanesia (SM cluster henceforth) and the other containing the remaining individuals (ANZO cluster). PC2 explained 6.8% of the variance and separated the two main archipelagos within SM: New Caledonia and Vanuatu. This population structure aligns with previous analyses using fewer individuals (Estandía et al. 2023). We visualised further substructure within the SM and ANZO clusters (Fig. S2 B-C).

We controlled for the population structure described above by using the eigenvectors that explained most of the variation, and conducted a GWAS that identified multiple loci linked to body size variation suggesting a polygenic basis for the trait. However, given the complicated population structure of our system, we retained those associated variants within the top 50 for which we had evidence of intra-population variation associated with body size, a total of 36 variants. We found that eight were classified as putative *de novo* mutations exclusive to islands in Southern Melanesia (Fig. 2A; Table S4), five of which were part of an association peak on Chromosome 2, which also contained one annotated gene, *GRHPR*, important in metabolic processes (Rumsby and Cregeen 1999). For the top outliers, those individuals with the genotype BB showed a five times increase in median body size relative to those with the AA genotype (Table S5), and the effect was still strong when we only explored Central Vanuatu, where the putative new mutation seems to be restricted to (Fig. 2A). This associated peak emerged again when we conducted GWAS on Southern Melanesia alone (Fig. S3). The effect size of the SNPs across this region was greater than any of the covariates included in the model to capture any population structure, suggesting no signs of conflation (Fig. S4). Other size-associated loci showed a similar effect in direction and magnitude on body size. Among these, were genes involved in calmodulin binding, cytoskeletal motor activity, organisation of the extracellular matrix and other metabolic processes (Table S6).

**Fig. 2.**
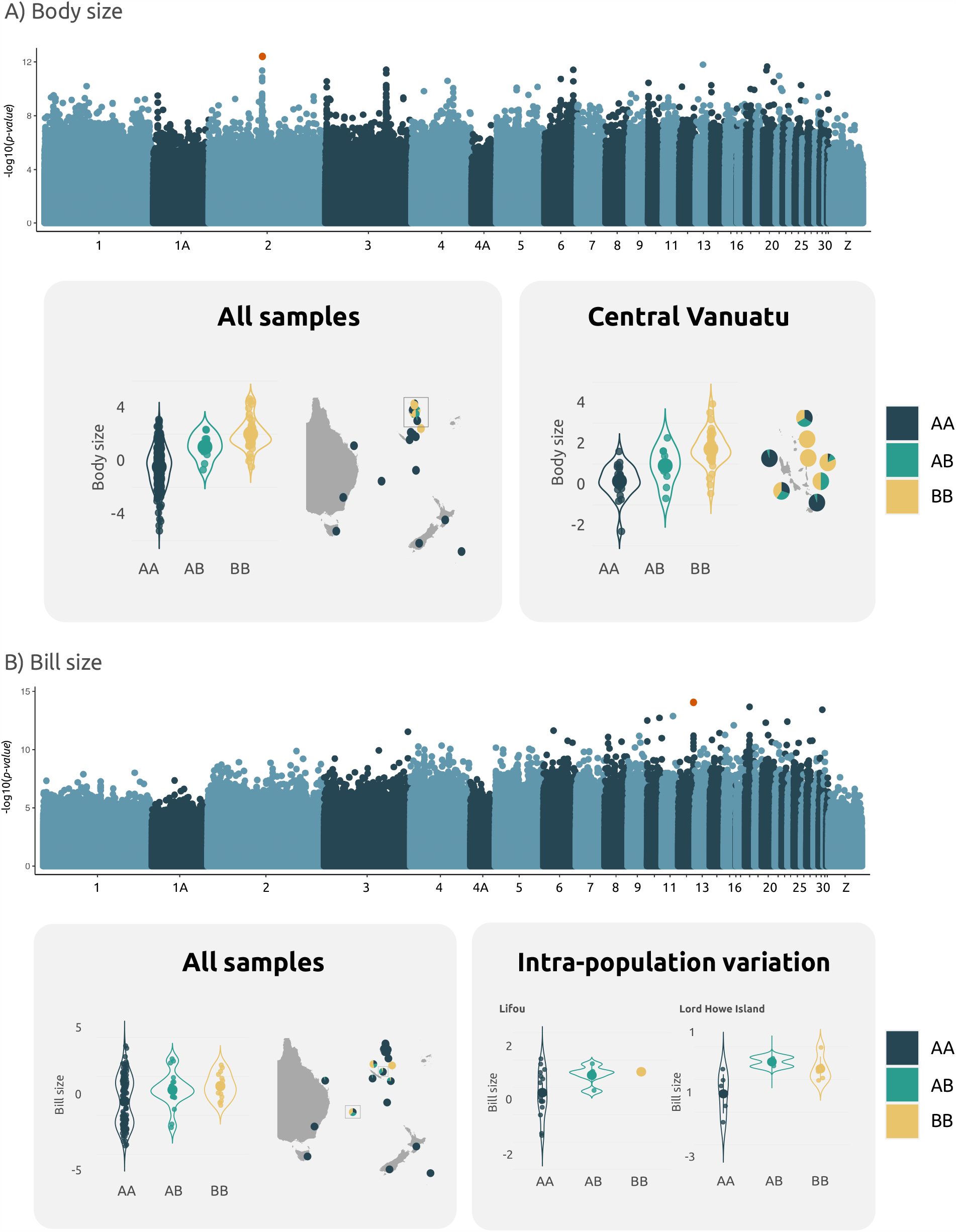
A) Manhattan plot for body size (PC1) and correlation between allele frequency for the top-ranked SNP and its geographic distribution across all samples (left) and restricted to Central and North Vanuatu, partly where the putative *de novo* mutations occur (right). It is important to note that this set of islands show minimal population structure. B) Manhattan plot for bill size (PC1) and correlation between allele frequency for the top-ranked SNP and its geographic distribution across all samples (left) and across two of the populations where there is intra-population variation associated with body size (right). These two populations are part of separate colonisations.

All of the top associated loci with bill size that were variable within populations represented standing genetic variation (Fig. 2B; Table S7). A number of adjacent variants on Chromosomes 12 and 18 were found in several of the large bill-size populations: islands in Southern Melanesia, Heron Island and Lord Howe Island. Among the annotated genes for bill size, we found gene *LGR6* (Table S8).

We quantified the proportion of phenotypic variance explained by all genome-wide SNPs with GCTA (Yang et al. 2011a). These narrow-sense heritability estimates were high but showed wide confidence intervals (CI) (Table S9). Tarsus length is a good proxy for body size and it is a highly heritable trait based on pedigree data, with a heritability of 0.7 for Heron Island silvereyes (Clegg et al. 2008), a higher value than our SNP-based estimate (h^2^=0.5 (95% CI =0.3-0.8)).

### Structural variation: a neo-sex chromosome and multiple putative inversions

To examine the structural variation along the genome, we identified clusters of PCA windows (local PCAs) based on Multidimensional Scaling (MDS) (Fig. 3-4). This method uses common patterns to identify structural variation such as non-recombining haplotype blocks (Li and Ralph 2019; Mérot et al. 2020). We found a large outlier region on Chromosome 4A (Fig. 3A-3B). MDS1 separated two distinct groups that aligned with sex (Fig. 3C), where sex was independently confirmed by PCR-based sexing of individuals.

**Fig. 3.**
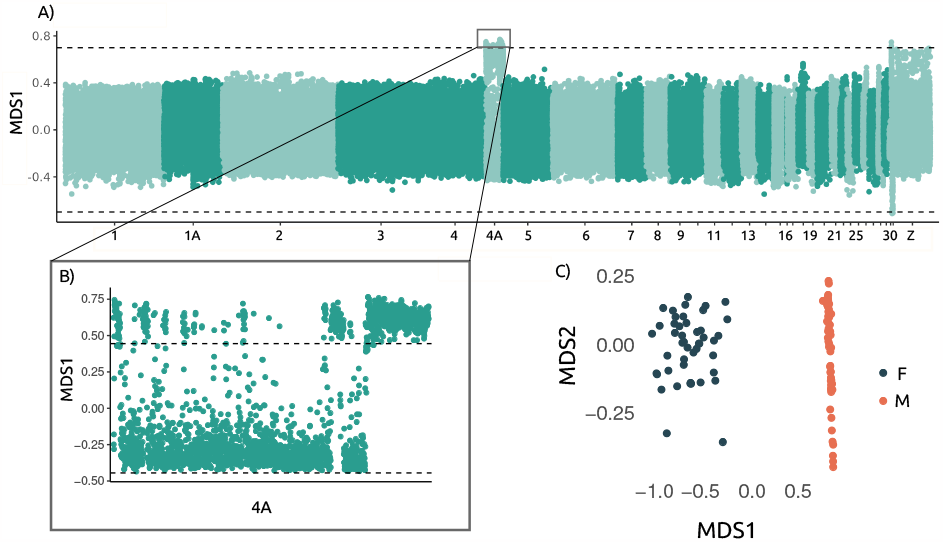
A) Local PCA results (MDS1) plotted across the genome B) where chromosome 4A shows outlier windows each representing 100 SNPs. C) MDS1 and MDS2 separate by sex in this region.

To specifically search for putative inversions, we filtered those sections of the genome with at least five adjacent outlier windows, pooled the windows and repeated MDS analysis for each separate region. Several chromosomes showed large outlier regions that formed three discrete equidistant clusters along MDS1. This type of pattern is highly suggestive of a potential inversion with each alternative homokaryotype separated by the intermediate heterokaryotype (Westram et al. 2022) (Fig. 4B). Consistent with the occurrence of inversions, LD was high inside these regions contrasting to the LD along the rest of the chromosome (Fig. S5), indicating limited recombination in heterokaryotypes. For the potential inversion on Chromosome 27, the AA karyotype (where A indicates the most common version of the arrangement) was present at high frequencies in all intermediate and old colonisations (Southern Melanesia, Lord Howe Island and Heron Island), and at very low frequency on the mainland. The potential inversion on Chromosome 28 showed a different pattern with the AA karyotype exclusively present in Southern Melanesia. On Chromosome 29, BB was only recovered from individuals in Southern Melanesia, but some individuals from New Caledonia displayed the AA karyotype (Fig. 4). Body size was not associated with karyotype for Chromosomes 27 and 29 after controlling for population structure. Body size was associated with the AA karyotype in Southern Melanesia, but we could not control for population structure as karyotype and neutral variation structure were entirely coincident. Using our custom annotation, we found genes involved in various metabolic processes on Chromosomes 27 and 28, and lipogenesis, adipocyte differentiation and muscle tissue development on Chromosome 29 (Table S10).

**Fig. 4.**
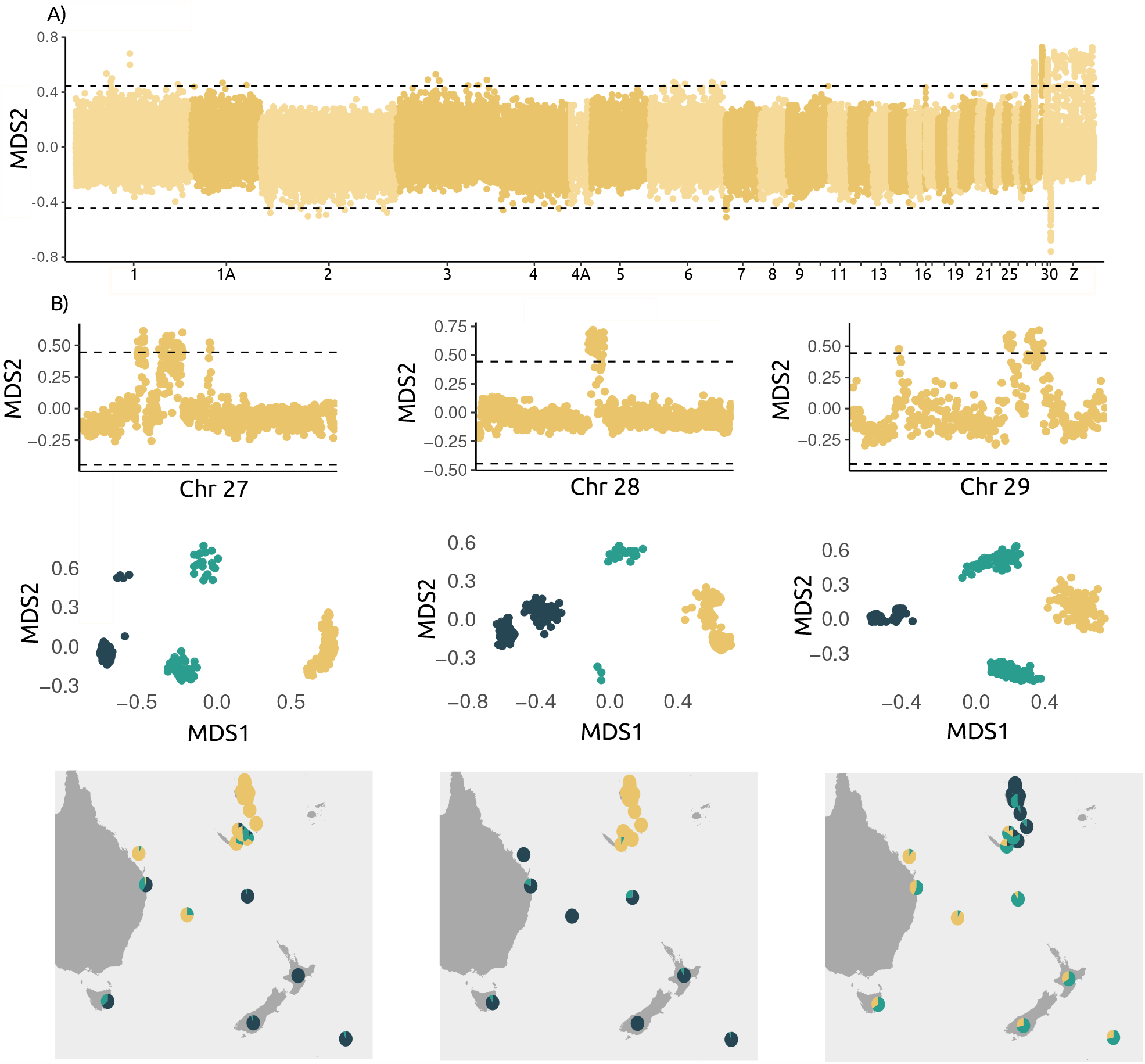
A) Local PCA results (MDS2) are displayed across the entire genome. B) Outlier regions in chromosomes 27, 28, and 29. Each dot corresponds to 100 SNPs. MDS1 identifies three distinct clusters within the outlier regions, suggesting potential inversions. Chartpies provide information about the relative frequencies of each cluster within different populations.

## Discussion

Selection on standing genetic variation is a common route to repeated evolution of larger body and bill sizes in island-colonising silvereyes. Size-associated variants were shared across colonisation histories spanning hundreds to over a hundred thousand years. However, putative *de novo* mutations present only in Southern Melanesia silvereye populations were also linked to increased body sizes. This empirical system suggests the joint role of standing genetic variation and *de novo* mutations in producing a repeated island phenotype.

We identified several candidate genes common across all populations. This pattern suggests that variation present in ancestral source populations was represented in multiple, independent colonising flocks, and maintained across sequential island colonisations. Given that population founding can result in loss of genetic variation and the fate of rare alleles can be highly stochastic especially when populations remain small (Whitlock 2000; Sendell-Price et al. 2021), such an outcome might seem unlikely. However, silvereyes colonise in reasonably large flocks (>100 individuals), and they recover rapidly from small population sizes, minimising loss of variation during founding and population establishment (Clegg et al. 2002a; Sendell-Price et al. 2021). Historically colonised silvereye populations and Heron Island silvereyes have experienced rapid trait evolution driven by natural selection (Clegg et al. 2002b, 2008; Sendell-Price et al. 2020), and such rapid evolution is more likely to stem from standing genetic variation than *de novo* mutations (Marques et al. 2019).

However, an association peak harbouring putative *de novo* mutations was present exclusively in Southern Melanesia, the first region to be colonised by silvereyes. These mutations were not fixed across all populations: some of them had fully fixed BB genotypes associated with very large body sizes, while others lacked the minor allele entirely, and many displayed all genotypes. This pattern held even within populations, indicating no bias due to population structure. Variation could be maintained across the archipelagos if different degrees of size are favoured in different populations connected by gene flow as seen across Southern Melanesian silvereye populations (Clegg and Phillimore 2010; Estandía et al. 2023). While larger size of island silvereyes appears generally to be favoured, other species, and indeed other *Zosterops* white-eye species that co-occur, could impose local constraints on size. For example, on Lifou, an island in New Caledonia, three Zosterops species occur in sympatry: the relatively small-sized endemic *Z. minutus*, the large endemic *Z. inornatus*, with the silvereye being intermediate in body size between the two (Mees 1969). An alternative but untested possibility is that variants arise from introgression with sympatric *Zosterops* species (Gill 1970). Other mechanisms for maintaining variation, such as negative frequency-dependent selection, are more difficult to reconcile in systems like islands where the usual drivers, e.g. predation risk, are limited (Beauchamp 2021).

Our study was unable to determine the prevalence of *de novo* mutations versus standing genetic variation. Even though we were able to detect new mutations in the oldest colonisation, our sample size was denser for this region than for any of the other of the old-to-intermediate colonisations which might have affected detection of association loci private to these populations. For these reasons, we can only say that in our study system, both routes seem to underlie body size variation, but their relative importance remains an important question to tackle in future studies.

Although linking annotated gene function to phenotypes in non-model organisms is highly speculative, any associations can provide the basis for future investigation. The putative *de novo* association peak on Chromosome 2 associated with body size has a hit for *GRHPR*. This gene encodes the enzyme glyoxylate reductase/hydroxypyruvate reductase, and is involved in key metabolic processes, including serine metabolism for gluconeogenesis and oxalate metabolism (Rumsby and Cregeen 1999). No other previous study has linked body size variation with this gene, yet, avian comparative genomic studies suggest that enzymes involved in glyoxylate metabolic routes might be responsible for diet-related adaptations in birds (Zhang et al. 2014). The function of other genes associated with body size is variable, from bone and muscle development to metabolic processing, and even though their role in body size is feasible, exact mechanisms remain to be found.

*LGR6* was one of the genes associated with bill size variation. This gene acts as a mediator for the Wnt/β-catenin signalling pathway, known to be involved in the formation of the avian beak (Bhullar et al. 2015). Interestingly, we did not find significant associations with *ALX1* or *HMGA2*, two large-effect genes that have been repeatedly demonstrated to be implicated in bill morphology variation, including in Darwin’s finches (Lamichhaney et al. 2015, 2016). However, different genetic architectures are not entirely surprising; even other species of finches show different pathways underlying bill size variation (vonHoldt et al. 2018).

We identified signatures of structural variants, one of which could potentially contribute to body size variation on chromosome 28, but others where such a relationship was clearly absent. Rearrangements on chromosomes 27, 28 and 29 each showed a pattern consistent with an inversion: three different clusters separated by MDS1 and high LD within the putative inversion compared with the rest of each chromosome (Mérot et al. 2020; Westram et al. 2022). Our conservative size estimates of the putative inversions observed here (approximately 200kb for those on Chromosomes 28 and 29 and 650 kb for Chromosome 27) place them in the range of inversion sizes known to be most frequently associated with adaptive phenotypes (100kb to several Mb) (Faria et al. 2019). Reduced recombination in the heterokaryotypes preserves linkage among alleles despite mating between alternative homokaryotypes, maintaining the adaptive benefit of the haplotype blocks (Faria et al. 2019; Westram et al. 2022). For example, marine snails (*Littorina saxatilis*) and seaweed flies (*Coelopa frigida*) show inverted regions linked to adaptive phenotypes (Morales et al. 2019; Koch et al. 2021); the latter has an inversion under balancing selection that works as a supergene for body size. The karyotype distribution of the putative inversion identified on Chromosome 28 was associated with body size but was conflated with geographic population structure. The function of genes inside this region varied widely, some of them are involved in immunity, others in cholesterol metabolism and some are even directly linked to fitness proxies in birds (Krist 2011), like egg quality (Gao et al. 2021). However, it is unclear whether they contributed to the evolution of large body sizes in Southern Melanesia. To further study the role of structural variants we could generate long-read sequencing and recombination maps that help to confirm that they are indeed inversions and to determine breakpoint locations (Mérot et al. 2020; Akopyan et al. 2022).

The three putative inversions displayed karyotype distribution patterns that suggested varying evolutionary histories. The inversion on Chromosome 28 could have emerged after colonisation from the Australian mainland and could have been subsequently fixed in the majority of Southern Melanesian populations. Aside from sampling considerations, alternative scenarios are more complex, requiring different combinations of karyotype loss and maintenance that seem less likely. For putative inversions on both Chromosomes 27 and 29, the maintenance of the heterokaryotype and alternative homokaryotype in many populations suggest they could represent ancient inversions under balancing selection. Such polymorphism can be maintained when the inversion provides a fitness advantage against different selective pressures: if there are epistatic interactions among alleles at different loci (Charlesworth and Charlesworth 1973), if there is associative overdominance (Ohta 1971), or if a locus that is under balancing selection is located within the inversion (Faria et al. 2019). Similar to alternative processes that could explain SNP variation, karyotypes could appear to be balanced if there is introgression resulting from gene flow among different sympatric Zosterops species occurring across Southern Melanesia, on Norfolk Island and prior to 20th-century extinction, also on Lord Howe Island, but not Heron Island or New Zealand.

A sizeable structural variant detected via local PCA and visualised by MDS was on chromosome 4A, coinciding with a known neo-sex chromosome in the Zosteropidae (Leroy et al. 2021), formed via fusion between an autosome and a sex chromosome approximately 24 Ma in the Sylvioidea superfamily to which the Zosteropidae belong (Pala et al. 2012; Sigeman et al. 2020). Neo-sex chromosomes can play a role in speciation as they can contain loci for traits that contribute to reproductive isolation (e.g. male courtship display behavioural traits (Kitano et al. 2009)). However, the role of this neo-sex chromosome in speciation within the Sylvioidea, including in the Zosteropidae, is unclear (Pala et al. 2012).

The study of non-model systems in the wild has inherent practical and bioinformatic limitations. Like any empirical study, we had to balance sample size and genomic coverage. GWAS usually relies on a high number of whole genomes with attached individual data. Capturing and taking samples from the birds used in this study covering a vast geographical distribution was a tremendous effort that spanned decades, and sequencing hundreds of genomes remains relatively expensive. Additionally, body and bill sizes are complex multivariate phenotypes that are highly polygenic (Duntsch et al. 2020) and correlate with several other traits (e.g. Estandía et al. 2021) that we did not investigate. Lastly, identifying the causal genes is a major challenge and often needs extra steps following GWAS to confirm the causality of the candidate loci, for example by doing knock-outs (Bengston et al. 2018), which is not possible in wild birds.

Despite these caveats and limitations, work on non-model systems is essential to gain a broad perspective on the different genetic routes that lead to similar phenotypes. The application of genomic methods to the silvereye system, which encompasses a wide range of island colonisation times and repeated evolution of body and bill size, provides a rare example of the role of standing genetic variation on both short and longer timescales of colonisation, with putative *de novo* mutations contributing only to the oldest island populations.

## Methods

### Sample collection

We sampled 377 silvereyes (*Z. lateralis*) from 31 islands or sites on the Australian mainland between 1997 and 2017 (Table S3). Birds were caught in the wild using mist nets or traps and we collected 20–40 μl of blood from the brachial wing vein and stored it in 1 ml of Queen’s lysis buffer (Seutin et al. 1991). We took a suite of morphological measurements, including the wing length using a metal ruler to determine the maximum flattened chord of the longest primary feather, and the length of the central tail feathers from the base to the tip using dividers. We used dial calipers (accuracy ±0.1 mm) to measure metatarsal length, head length from the rear of the skull to the bill’s tip. Additionally, we took various bill measurements using dial calipers, including mandible length, width, and depth at the posterior nostril opening, and mandible length, width, and depth at the anterior nostril opening. Finally, we measured the body weight. Sex cannot be determined in silvereyes with morphological measurements. We selected a random subset of individuals and sexed them using molecular methods described in the section below. All sampling was non-destructive, and we released the birds at point of capture. Complete sampling information is provided in Table S3.

### Construction of a pseudochromosome assembly

We used the Chromosembler tool available in Satsuma2 (Grabherr et al. 2010) to order and orient scaffolds of the Z. l. melanops genome assembly (GCA_001281735.1) (Cornetti et al. 2015) into 33 pseudo-chromosomes according to synteny with the Vertebrate Genome Project’s Zebra finch (*Taeniopygia guttata*) genome assembly (GCA_003957565.1) (Rhie et al. 2021).

To reconstruct a transcriptome that we could use to generate hints for gene prediction, we used available RNA-seq data for *Zosterops borbonicus* (Leroy et al. 2021). We filtered out low-quality sequence reads (phred quality score < 20) and adaptor sequences using TrimGalore (Martin 2011) and mapped them onto our pseudo-chromosome assembly using HISAT2 (Kim et al. 2015). We used Trinity (Grabherr et al. 2011) to assemble the transcriptome.

### Assembly annotation

We annotated the resulting pseudo-chromosome assembly using a multifaceted approach of repeat identification and *ab initio* gene prediction using AUGUSTUS (Stanke et al. 2006). We incorporated hints containing extrinsic evidence about the location and structure of three types of genomic features: repetitive regions, short-read transcriptome sequences, and single-copy orthologs. To obtain our hints, we first identified and masked repetitive and low-complexity DNA sequences using RepeatMasker (Smit et al. 2013) with the chicken (*Gallus gallus*) database of repeats. Second, we used BLAT (Kent 2002) to incorporate RNA-seq information as hints. Third, we used single-copy genes in the Aves lineage group odb10 to detect conserved orthologous genes with BUSCO (Manni et al. 2021). We then used BLASTx (Camacho et al. 2009) in the AUGUSTUS output to obtain functional information. More information about specific parameters used, including cut-offs, can be found at https://github.com/andreaestandia/1.0_island_rule_GWAS/tree/main/pipeline/9_annotation

### DNA extraction, molecular sexing, library preparation, and sequencing

We extracted DNA from blood samples using a standard Phenol-chloroform protocol (Seutin et al. 1991). We added 100 uL of blood stored in Queen’s buffer to a microcentrifuge tube containing 250 uL of DIGSOL extraction buffer (0.02 M EDTA, 0.05 M Tris-HCl (pH 8.0), 0.4 M NaCl, 0.5% sodium dodecyl sulphate (SDS)) and 10uL of Proteinase K (20mg/mL). Samples were incubated at 55C overnight. Following incubation, we added 250 uL of phenol:chloroform:isoamyl alcohol (25:24:1) and gently mixed the samples for 10 minutes. We then centrifuged the samples at 10,000 rpm for 10 minutes. We transferred the aqueous layer to a new microcentrifuge tube and repeated the previous step by adding the phenol mixture, mixing and centrifuging it. We recovered a new aqueous phase, transferred it to a new tube, added 250uL of chloroform:isolamyl alcohol (24:1), mixed and centrifuged the samples as in the previous steps. We precipitated the DNA by adding 2 volumes of ice-cold 100% ethanol, 1 volume of 2.5M ammonium acetate and 2 uL of glycogen, leaving it overnight at -20C. The next morning we centrifuged the tubes for 10 min at 15,000 rpm at 4C and removed the supernatant. We rinsed the precipitate with 500 uL of ice-cold 70% ethanol and centrifuged the tubes as in the previous step. We removed the supernantant and the left the precipitated DNA to dry at room temperature. Once dried, the DNA was resuspended in 50 uL of TE (TrisEDTA) buffer (0.01 M Tris-HCL (pH 8.0), 0.0001 M EDTA). We quantified the DNA concentration with a Qubit 2.0. DNA extracts were sent to Novogene UK for library preparation and whole genome sequencing at 5X on the Illumina Novaseq 6000 platform (Illumina, San Diego) using paired-end 150bp sequencing reads.

### Quality control and genotype calling

We trimmed reads to remove adapter content and base calls of low quality using fastp (Chen et al. 2018) with default settings and 10 bp trimmed from the start of each read. We aligned cleaned reads of each individual to our pseudochromosome assembly using the Burrows-Wheeler Aligner (BWA)-mem algorithm (Li and Durbin 2010). Genotype calling can result in bias when using low-coverage sequencing (Johnson and Slatkin 2007). Given the moderately low sequencing depth of our dataset (5X) downstream analyses were based on genotype probabilities rather than directly-called genotypes. We calculated genotype probabilities in ANGSD (Korneliussen et al. 2014) using the samtools model and keeping SNPs with a minimum allele frequency (MAF) of 0.05. All analyses described below were run considering only biallelic SNPs with a mapping quality score 20. We filtered the bam files by excluding SNPs that showed significant deviation from Hardy-Weinberg equilibrium, recovering a total of 14,330,920 SNPs.

### Statistical tests of repeated increase in body and bill size

Body size is most often represented by multivariate measures, such as Principal Component Analysis (PCA), that account for the covariation among characters and summarise most of the variation in the first two axes, the first representing body size and the second body shape (Rising and Somers 1989). We conducted a PCA that included all morphological measurements that represent body size: wing length, tail length, tarsus length, and head length; and separately bill size: bill length to posterior nostril opening, bill width, and depth at anterior nostril opening. We tested whether colonisations resulted in changes in body size and bill size relative to the mainland using *brms* (Bürkner 2017). Model 1 included PC1 of body size as the response variable and Model 2 PC1 included bill size. Colonisation time was set as a categorical population-level effect with a weak prior in the form of a normal distribution centered in 0 with a standard deviation of 10. This prior gives equal support to the idea that body size can remain unchanged, increase or decrease after colonisation. We controlled for population by setting it as a group-level effect. We used a Gaussian family and the default link function. We ran four chains each with 5000 iterations which included a warm-up of 1000.

### The genetic basis of parallel evolution

To test whether *de novo* mutations or ancestral standing genetic variation led to similar phenotypes, we did a Genomewide Association Study (GWAS) under the idea that if the associated variants are common to all populations then they are likely due to standing genetic variation but if they are present in a specific population or specific lineage, they may represent *de novo* mutations that have accumulated since the lineage split. This framework assumes no potential gene flow among colonisations, which is supported by genome-wide data (Estandía et al. 2023).

#### Genome-wide Association Study

We conducted GWAS on the first two PCs of body size (body size PC1 and body size PC2) and bill size (bill size PC1 and bill size PC2). We ran a generalised linear model implemented in PLINK (Purcell et al. 2007) and controlled for population structure by generating a covariance matrix with PCAngsd (Meisner and Albrechtsen 2018) and decomposing it by using a scaling transformation. We used the first 8 eigenvectors that accounted for most of the variation and adding more components accounted for less than 1% of the variation and did not split populations further. The two main clusters (ANZO and SM) have diverged for a long time, and many of the associated variants could have arisen due to population substructure not represented by the top eigenvectors and phenotypes that did not fully overlap between the ANZO and SM groups (but see Fig. S6). To check that our results held on each subset, we repeated the GWAS on each subgroup while accounting for their own population substructure (Fig S2 B-C). For the top 50 associated variants, we excluded samples with genotype likelihoods lower or equal to 0.33, which indicates that all genotypes have the same probability of being the true genotype. We classified putative *de novo* mutations as those only present in a single colonisation. For example, if an associated variant is only present on Heron Island, it would be classified as a new mutation. However, if the associated variant is present in various populations from separate colonisations, we classified them as arising due to standing genetic variation. This is because it is unlikely that the same new mutation would arise more than once in separate lineages stemming from the same source. Given that our system shows a complicated population structure, we decided to be conservative and further filtered associated variants by keeping those that showed at least one population where there was evidence of intra-population variability associated with body size. We did this by conducting an ANOVA between genotype and body size in each population separately. We also explored the frequency of genotypes and plotted it on maps to explore geographic structuring. We used our custom annotation to find the regions where the outliers were located by selecting all annotated coding regions within 5000 bp upstream and downstream from the associated locus. To confirm that our annotation was robust, we pulled the same number of bp from the reference genome and performed a BLAST search against all organisms. To estimate the proportion of phenotypic variance explained by all genome-wide SNPs while correcting for LD bias we used GCTA (Yang et al. 2011a). We created four quartiles of LD scores and used each individual SNP’s LD score for binning as recommended by Evans et al. 2018. We also controlled for population structure using the principal components arising from the PCAngsd analysis. We computed the h^2^-SNP 95% CI as +-1.96 * SE.

### Detection of structural variants

To detect structural variants, we conducted a localised PCA of genetic variation based on the methodology outlined in Li and Ralph (2019). We used PCAngsd and the R package *lostruct* to identify non-recombining haploblocks across the genome. Structural variants like inversions are characterised by high levels of linkage disequilibrium because recombination is often to be negligible in heterokaryotes (Kirkpatrick 2010). For this reason, we used the unpruned SNP dataset containing 14 million SNPs. First, we split each chromosome into non-overlapping windows containing 100 SNPs each, ran PCAngsd and obtained a covariance matrix for each window. We then descomposed each matrix by using scaling transformation and generated a pairwise distance matrix based on PC1 and PC2 for each window. Finally, we mapped similarity using MDS of up to 20 axes. We defined outlier regions as windows with a coefficient of correlation above four standard deviations from the mean. Adjacent outliers with less than 10 windows between them were pooled, and clusters with less than 5 windows were not considered following Mérot et al. (2021). A typical signature of a polymorphic inversion is three groups of individuals appearing on a PCA: the two homokaryotypes for the alternative arrangements and, as an intermediate group, the heterokaryotypes (Mérot et al. 2020). All clusters of outlier windows were examined by MDS as single blocks. We then used K-means clustering to identify putative groups of haplotypes. We explored LD patterns in heterokaryotype versus homokaryotype individuals for the chromosomes where we found outlier regions. We calculated pairwise r^2^ values with ngsLD based on genotype likelihoods obtained by ANGSD and created a heatmap where we could visually explore LD levels. We built a model in *brms* to test whether karyotype identity was associated with body size after controlling for population structure. Karyotype was set as a categorical population-level effect and body size as the explanatory variable. Population was set as a group-level effect. We used a Gaussian family and the default link function. We ran four chains each with 5000 iterations which included a warm-up of 1000. We refer to karyotypes AA, AB, and BB where A represents the most frequent variant, whether inverted or not since we do not have information about this.

## ACKNOWLEDGEMENTS

The samples used in this study were collected over two decades and we thank the many people who helped in numerous ways to facilitate the work. We are grateful to the chiefs and landholders of Vanuatu and New Caledonia for granting access to field sites, and field and logistic assistance from many people including the following: N. Clark, D. Treby, J. LeBreton, F. Cugny, W. Waheoneme, O. Hebert and A. Rouquie (New Caledonia); F. Robertson and C. Sendell-Price (Heron Island); S. Geiger, O. Boissier, R. Hills, S. Totterman, O. Drew, K. Ser, W. Ser, J. Saksak (Vanuatu), I.P.F. Owens, N. Clark (Australian mainland); J. Kikkawa (Norfolk Island); I.P.F. Owens (Lord Howe Island); P. Park, A. Fletcher, P. Gray (Tasmania); P. Schweigman, D. Onely (New Zealand); M. Bell (Chatham Island); N. Davies, O. Grant, C. Sendell-Price (Moorea, French Polynesia). For additional samples we thank A. Phillimore, R. Black, M. Massaro, D. Potvin and N. Clark. The work was conducted under permits from the Direction de l’Environnement Province Sud and Direction Du Développement Economique (New Caledonia and Loyalty Islands with thanks to G. Kakue); Vanuatu Environment Unit letters of permission and permits provided by E. Bani and we further thank D. Kalfatak and T. Tiwok for their assistance (Vanuatu); Délégations régionales à la recherche et à la technologie (French Polynesia); Lord Howe Island Board; Environment Australia (Norfolk Island); Queensland Department of Environment and Resource Management; Parks and Wildlife Service (Tasmania). New Zealand Department of Conservation Te Papa Atawhai; Australian Bird and Bat Banding Scheme project and individual permits to SMC. Ethics clearances were provided by University of Queensland ethics committee (ZOO/165/94/ARC, ZOO/520/96/ARC/PHD, ZOO/520/97/ARC/PHD) and Griffith University ethics committee (ENV/01/12/AEC, ENV/07/16/AEC, ENV/06/20/AEC, ENV/24/13/AEC) to SMC. We thank F. Robertson for conducting labwork. We thank the funders of this work: a Marsden Fund grant (UOO1410) from the New Zealand Royal Society Te Apārangi awarded to BCR and SMC to support fieldwork on mainland Australia and Heron Island portions of the molecular work; National Geographic Society Committee for Research and Exploration Grant (9383-13) to SMC to support fieldwork in New Caledonia; Natural Environment Research Council (NERC) postdoctoral fellowship to SMC to support fieldwork in Vanuatu and New Caledonia; Percy Sladen Memorial fund to SMC to support fieldwork in French Polynesia; the Department of Zoology, Oxford startup to SMC to support sequencing. ASP was supported by a NERC studentship (NE/L002612/1); AE was supported by a NERC studentship (NE/S007474/1) and a St John’s College Graduate Scholarship.

## Author contributions

SMC and AE formulated the idea and SMC supervised the project. SMC, BCR, and ATSP conducted fieldwork. AE, ATSP and BCR conducted laboratory work. AE conducted the genetic and statistical analyses with input from ATSP. AE drafted the manuscript with input from SMC and all authors provided comments.

## Data and code availability

Code to reproduce all analyses is available at https://github.com/andreaestandia/1.0_island_rule_GWAS

