## Supplementary figures and images for "Standing genetic variation and *de novo* mutations underlie parallel evolution of island bird phenotypes"

### Fig. S1

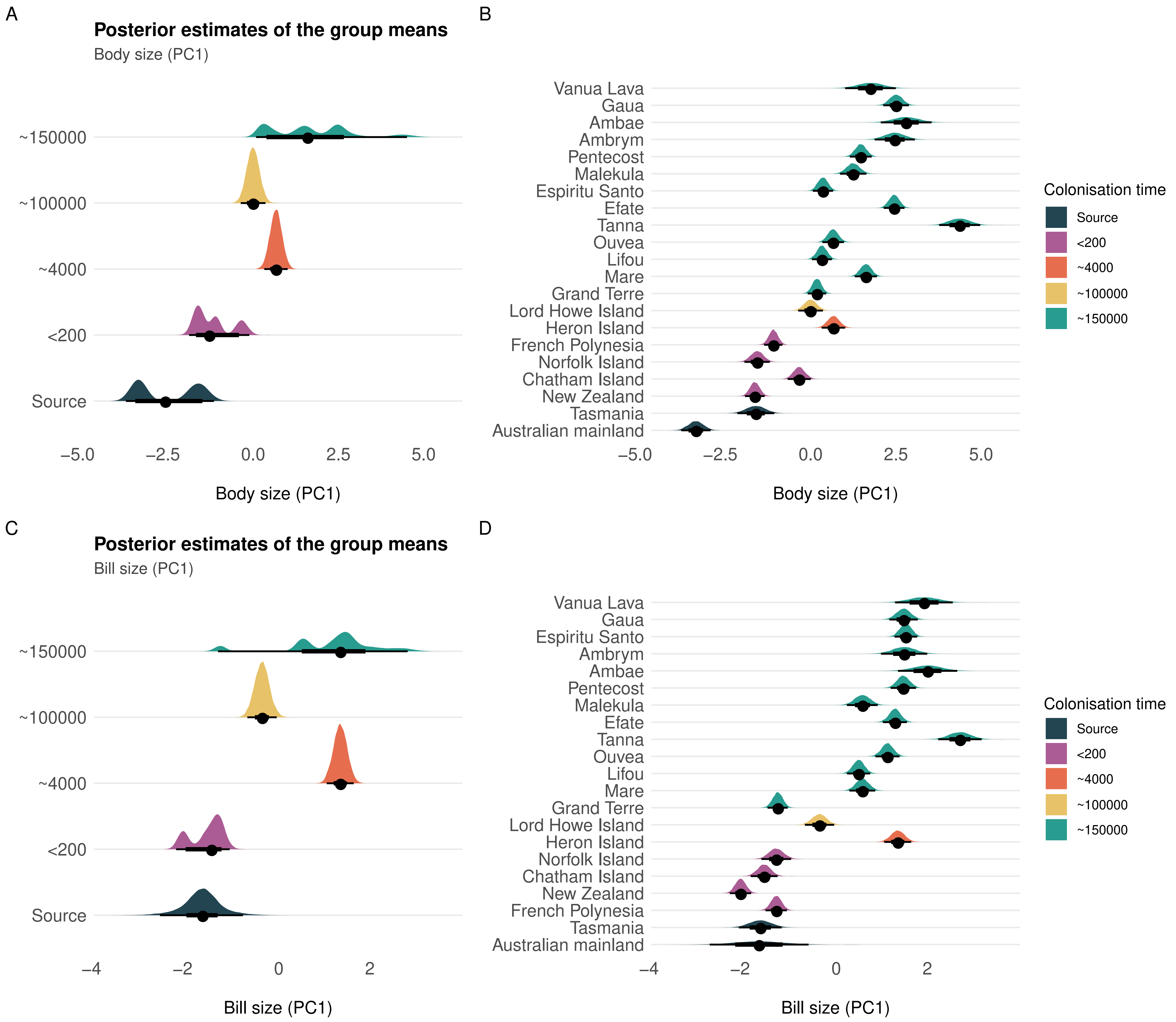

### Fig. S2

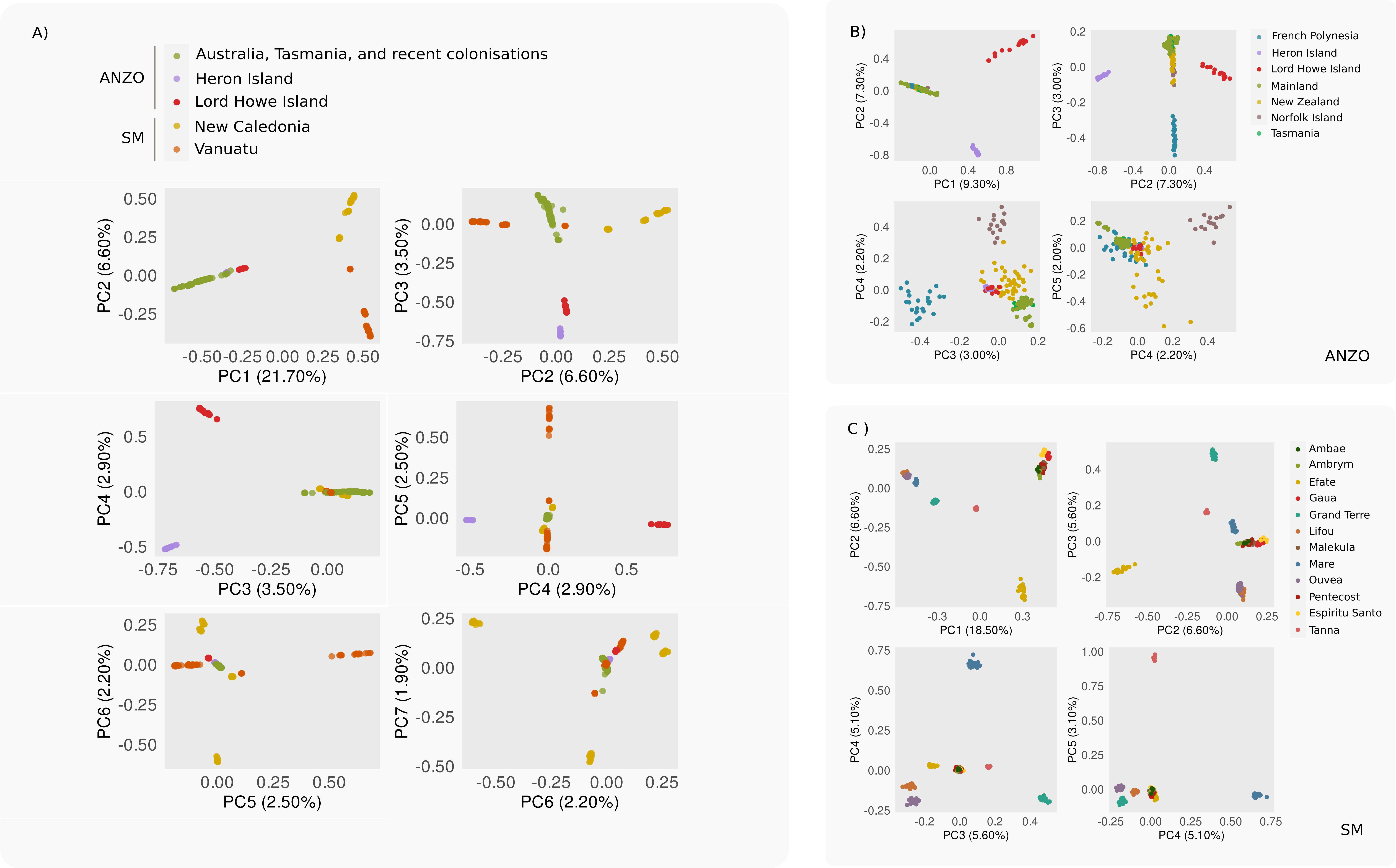

### Fig. S3

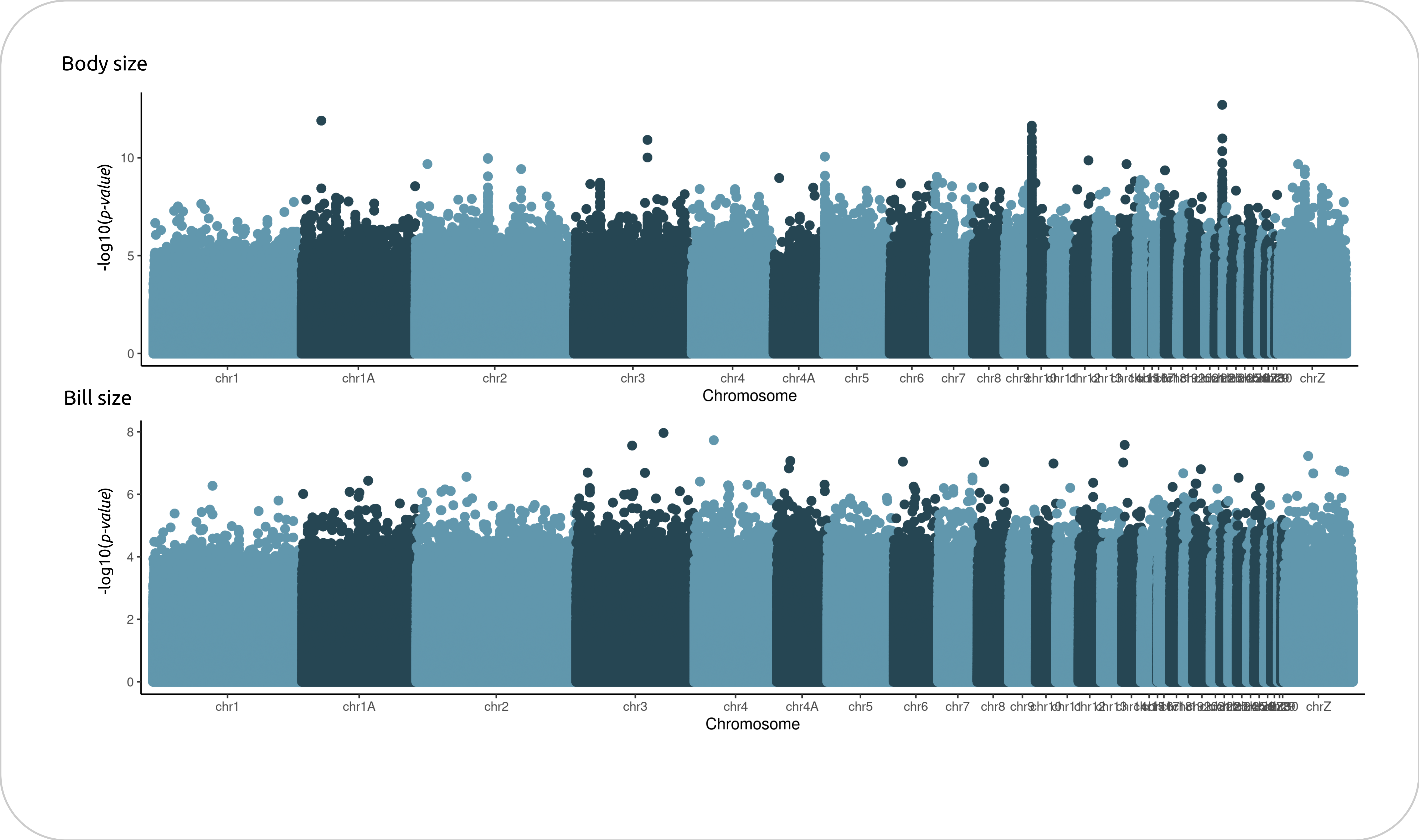

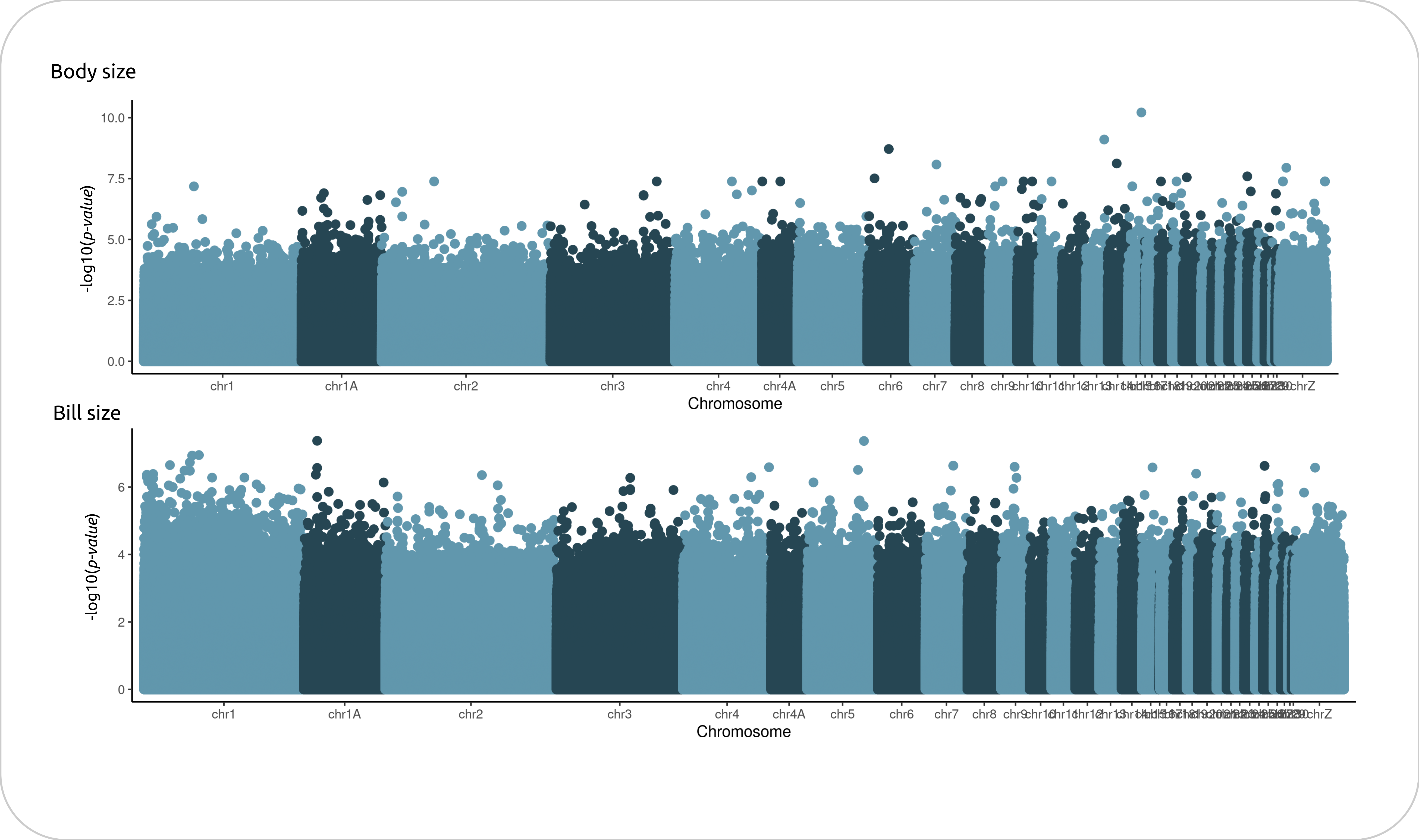

### Fig. S4

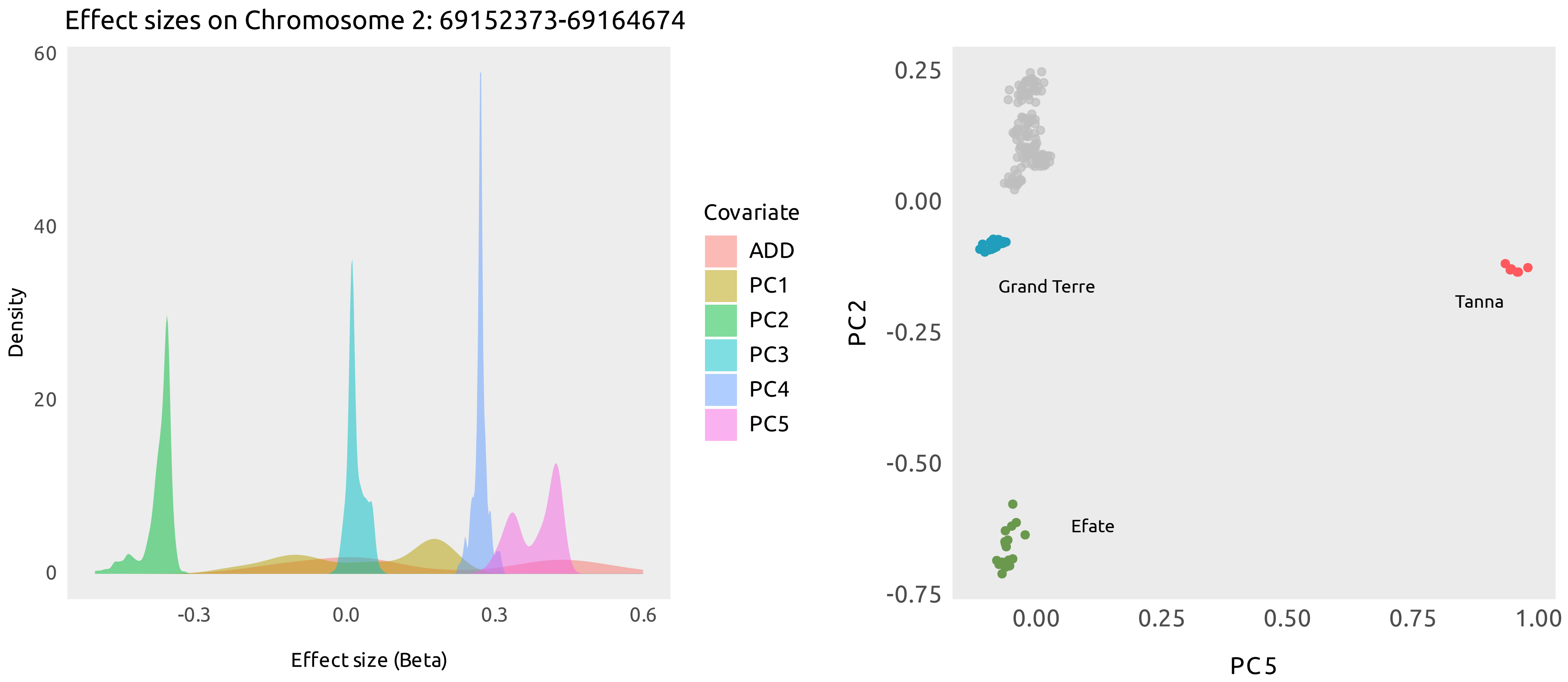

### Fig. S5

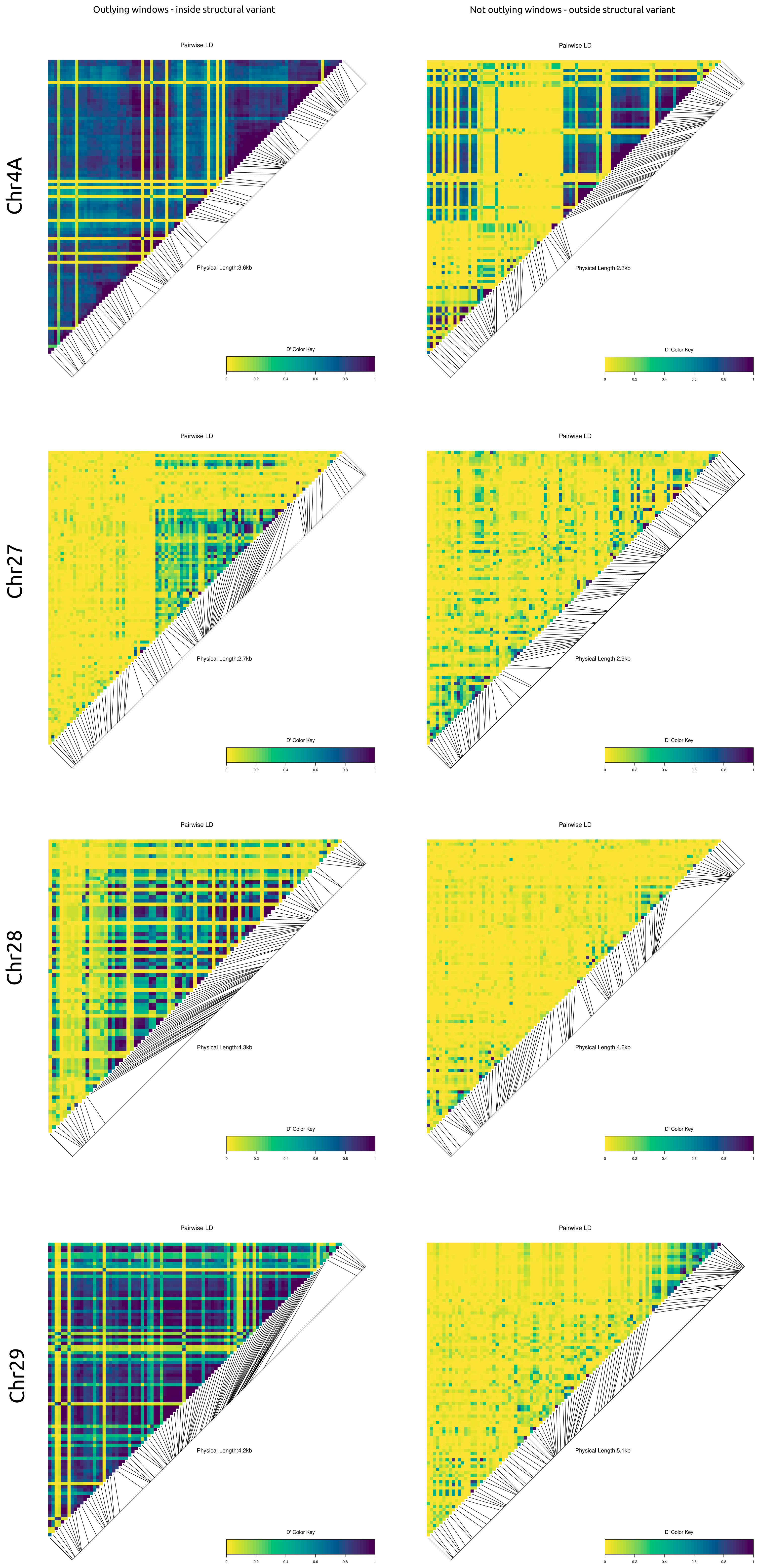

### Fig. S6

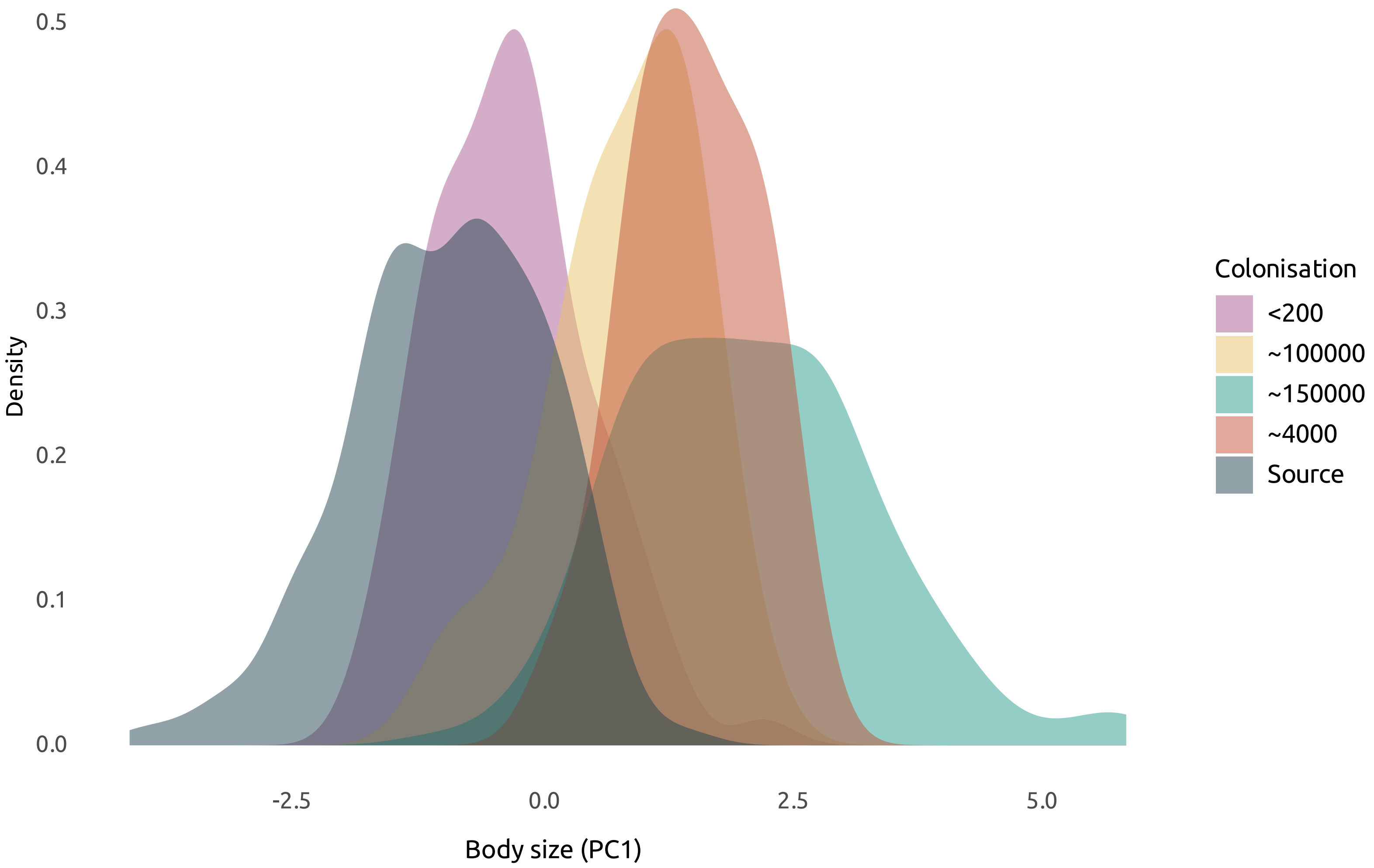
